# First description of Hepatitis E virus in Australian rabbits

**DOI:** 10.1101/2020.10.05.327353

**Authors:** Maria Jenckel, Robyn N. Hall, Tanja Strive

## Abstract

We report the first detection of Hepatitis E virus in rabbits in Australia. While conducting metatranscriptomic sequencing of liver samples collected from domestic rabbits that had diedwe detected three samples positive for hepatitis E virus. Two viral genome sequences were obtained, which shared 96% nucleotide identity.

---

Mammalian hepatitis E viruses (HEV) are non-enveloped positive-sense monopartite RNA viruses in the family *Hepeviridae*, genus *Orthohepevirus*, species *Orthohepevirus A*. Viral genomes are approximately 7200 nt in length and consist of three open reading frames (ORFs). The non-structural proteins are encoded in ORF1, whereas ORF2 codes for the capsid protein and ORF3 for a small phosphoprotein (1). HEV are genetically diverse and infect a wide range of vertebrate species, with four genogroups (HEV-1-4) reported to infect humans. The primary route of infection is fecal-oral, and contact with infected animals, contaminated water sources or contaminated food products, especially undercooked pork, are considered to be the main sources of human infections (2). HEV infections are usually asymptomatic and self-limiting, however, immunocompromised patients and pregnant women can develop much more severe disease, with case fatality rates of up to 25% (2).

HEV was first reported in farmed rabbits (*Oryctolagus cuniculus*) in China in 2009 (3) and was assigned to genotype 3 (HEV-3) based on phylogenetic analyses. However, retrospective serological analyses showed that the virus was present in wild rabbits in Europe in the late 1980s (4). In 2016, HEV-3 viruses closely related to those found in rabbits were isolated from human patients, suggesting zoonotic transmission (5). Infection in rabbits appears to be asymptomatic, although subclinical hepatitis may be induced (6).

As part of a study aimed at better understanding the pathogens present in Australian rabbits, we analysed 48 liver samples from domestic rabbits that had died between March 2017 and July 2019 using a metatranscriptomics approach. Total RNA was extracted from liver samples and libraries were generated using the NEBNext® Ultra™ II RNA Library Preparation Kit (New England Biolabs) including an rRNA depletion step (NEBNext® rRNA Depletion, New England Biolabs). Samples were sequenced on an Illumina NovaSeq6000 instrument using an SP300 cycle flow cell. Adapters were trimmed and low-quality reads were discarded. Host sequences were removed by mapping data to the rabbit reference genome (GCF_000003625.3) using bowtie2. The remaining reads were assembled using Trinity and contigs were checked against the NCBI nucleotide database. Contigs and reads assigned to HEV were assembled into full-length or partial viral genomes using SPAdes.

HEV-3 was detected in three of the 48 sequenced samples. One was from a 3-year-old male Rex rabbit (DDS-1) admitted to a shelter and subsequently euthanised for suspected myxomatosis in November 2018. Clinical signs included swelling of the eyelids and scrotum and tachypnoea. On post-mortem examination there was consolidation of the lungs, hepatomegaly, and blood in the abdomen. The second detection (CGW-40) was from a juvenile mini lop rabbit that died in a veterinary hospital in January 2019 with hemothorax present on post-mortem examination. The third detection was from a 10-week-old male mini Rex rabbit (HTL-10) that appeared bloated and lethargic and subsequently died, along with four other rabbits from a litter of six, in July 2019. All three samples were collected from domestic rabbits located in Victoria, in south-eastern Australia.

The complete HEV genome was recovered from HTL-10 (Genbank MW002523), along with a near complete genome from DDS-1 (Genbank MW002522) missing the first 70 nucleotides. Thirty-six sequencing reads were obtained from CGW-40, and positivity was further confirmed by RT-qPCR (7).

BLASTn results of the two Australian HEV-3 genomes showed 96% nucleotide identity to each other and 87% nucleotide identity to a human HEV strain from France collected in 2008 (JQ013793). This French virus clusters with rabbit HEV-3 strains identified in wild and farmed rabbits from France (2007-2010) (JQ013791, JQ013792) and farmed rabbits from China in (2009-2010) (FJ906895, FJ906896, GU937805) (5). The closest matches to HEV-3 sequences specifically from rabbits showed nucleotide identities of 85.2% (KX227751, China, domestic rabbit, 2015) and 85.8% (JX565469, USA, domestic rabbit, 2010) for HTL-10 and DDS-1, respectively. Both Australian HEV-3 sequences harbor a 93 nt insertion in the X domain of ORF1, common to all rabbit HEV-3 strains (8). Amino acid identities ranged from 93.4 – 97% for protein-coding regions (Table 1).

**Table 1:** Amino acid identities for ORF 1-3 of Australian HEV-3 strains

|  | HTL-10 |  | DDS-1 |  |
| --- | --- | --- | --- | --- |
|  | Closest amino acid identity | Genbank Accession | Closest amino acid identity | Genbank Accession |
| ORF 1 | 94.2 % | JQ013793 | 93.4 % | AB740221 |
| ORF 2 | 96.8 % | KY436898 | 97 % | KX227751 |
| ORF 3 | 95.1 % | MF480297 | 94.3 % | MF480297 |

Phylogenetic analysis of the HEV-3 genomes recovered in this study, along with representative published sequences, shows that the generated sequences clearly formed a monophyletic clade with other rabbit HEV-3 strains (Figure 1).

**Figure 1.**
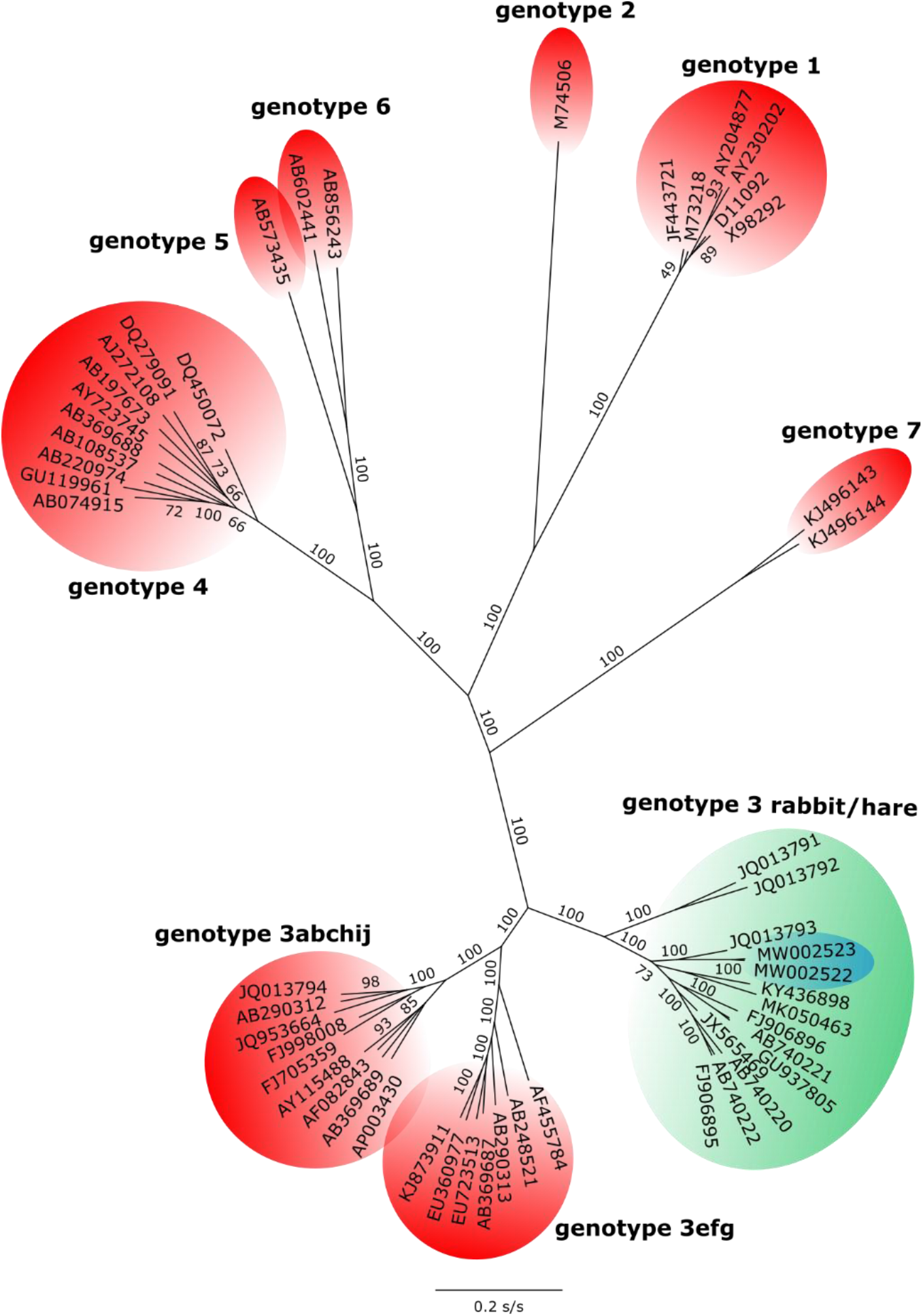
An unrooted maximum likelihood tree of representative HEV genomes was constructed using IQ-TREE (v. 2.0.6) with the best fitted model (GTR+F+R5) and 1000 ultra-fast bootstraps. Reference genomes and genogroups were selected based on Smith et al. (9). Genotype 3 (rabbit and hare) sequences are highlighted in green. New sequences from this study are highlighted in blue. Bootstrap values are shown for selected nodes. The scalebar shows substitutions per site (s/s).

Despite the liver samples in this study being collected from sick animals, there is no clear evidence that HEV-3 infection was a contributing factor in the death of these rabbits. All samples analysed in this study were obtained from domestic animals and it is not yet known whether wild rabbits and hares in Australia also carry HEV-3. Due to the increasing popularity of domestic rabbits as pets, as well as the abundance of wild rabbits in Australia, the detection of HEV strains that are closely related to known zoonotic HEV-3 strains raises concerns for potential zoonotic transmission to humans. More comprehensive analyses are needed to determine the geographical distribution and host range of HEV-3 in Australia.

## Acknowledgements

The project was co-funded by Meat and Livestock Australia (P.PSH.1059) and CSIRO. Rabbit samples were obtained through a project funded by the Centre for Invasive Species Solutions (P01-B-002).

## References

1. Purdy MA, Harrison TJ, Jameel S, Meng XJ, Okamoto H, Van der Poel WHM, et al. ICTV Virus Taxonomy Profile: Hepeviridae. J Gen Virol. 2017 Nov;98(11):2645–6.

2. Nan Y, Wu C, Zhao Q, Zhou EM. Zoonotic Hepatitis E Virus: An Ignored Risk for Public Health. Front Microbiol. 2017;8:2396.

3. Zhao C, Ma Z, Harrison TJ, Feng R, Zhang C, Qiao Z, et al. A novel genotype of hepatitis E virus prevalent among farmed rabbits in China. J Med Virol. 2009 Aug;81(8):1371–9.

4. Eiden M, Vina-Rodriguez A, Schlosser J, Schirrmeier H, Groschup MH. Detection of Hepatitis E Virus in Archived Rabbit Serum Samples, Germany 1989. Food and Environmental Virology. 2015;8(1):105–7.

5. Izopet J, Dubois M, Bertagnoli S, Lhomme S, Marchandeau S, Boucher S, et al. Hepatitis E virus strains in rabbits and evidence of a closely related strain in humans, France. Emerg Infect Dis. 2012 Aug;18(8):1274–81.

6. Li S, Li M, He Q, Liang Z, Shu J, Wang L, et al. Characterization of hepatitis E virus natural infection in farmed rabbits. J Viral Hepat. 2020 Aug 27.

7. Jothikumar N, Cromeans TL, Robertson BH, Meng XJ, Hill VR. A broadly reactive one-step real-time RT-PCR assay for rapid and sensitive detection of hepatitis E virus. J Virol Methods. 2006 Jan;131(1):65–71.

8. Wang L, Liu L, Wang L. An overview: Rabbit hepatitis E virus (HEV) and rabbit providing an animal model for HEV study. Rev Med Virol. 2018 Jan;28(1).

9. Smith DB, Simmonds P, Izopet J, Oliveira-Filho EF, Ulrich RG, Johne R, et al. Proposed reference sequences for hepatitis E virus subtypes. The Journal of general virology. 2016;97(3):537–42.

